# Selective Smurf1 E3 ligase inhibitors that prevent transthiolation

**DOI:** 10.1101/2023.10.14.562361

**Authors:** Patrick Chuong, Alexander Statsyuk

**Author notes:** **Corresponding Author** Alexander Statsyuk – Department of Pharmacological and Pharmaceutical Sciences, College of Pharmacy, University of Houston, Houston, TX 77004, United States.

## Abstract

Smurf1 is a HECT E3 ligase that is genetically micro-duplicated in human patients and is associated with osteoporosis. Smurf1^-/-^ mice on the other hand show an increase in bone density as they age, while being viable and fertile. Therefore, Smurf1 is a promising drug target to treat osteoporosis. This paper reports the discovery, synthesis, and biochemical characterization of highly selective Smurf1 inhibitors. We show that these compounds inhibit the catalytic HECT domain of Smurf1 with 500 nM IC_50_, but they do not inhibit closely related Smurf2 ligase, which is 80% identical to Smurf1. We show that Smurf1 inhibitors act by preventing the trans-thiolation reaction between Smurf1 and E2∼Ub thioesters. Our preliminary studies show that the C-lobe of Smurf1 alone does not contribute to the observed high selectivity of Smurf1 inhibitors.

---

HECT E3 ligases are a small family of E3 ligases (∼28 known) that range in size from 100 kDa to 500 kDa and have a variety of physiological functions.[1] Genetic mutations of HECT E3s in humans or deletion in mice are associated with the variety diseases and phenotypes.[1]–[4] Understanding physiological functions of HECT E3s would be greatly facilitated by pharmacological inhibitors of these enzymes that can also be used to identify cellular substrates of HECT E3s using quantitative proteomic methods.[5] However, as of today, highly selective and well characterized inhibitors of HECT E3 ligases are not known. Smurf1 is a HECT E3 ligase that regulates bone growth in mice and humans.[6]–[8] It is believed that Smurf1 regulates bone growth by inhibiting the function of osteoblasts, the cells that produce the bone matrix. Some of known Smurf1 substrates in osteoblasts include transcription factor RunX2, kinase MEKK4, and <u>ins</u>ulin receptor (InsR).[6], [7] Degradation of RunX2 and MEKK4 by Smurf1 are believed to inhibit osteoblast function, while degradation of InsR by Smurf1 is responsible for the observed hypoglycemia in Smurf1^-/-^ mice.[7] Smurf1 inhibitors that target WW domains of Smurf1 have been reported, however inhibitors that target the catalytic HECT domain are not known. [9] This paper reports the characterization of highly selective Smurf1 inhibitors and their mechanistic studies.

Searching the patent literature revealed a group of Smurf1 inhibitors with pyrazolone core motifs that, according to the patent, displayed single digit nanomolar IC_50_’s against Smurf1, and showed no inhibition of the closely related Smurf2 HECT E3.[10], [11] The lack of inhibition of Smurf2 is remarkable because Smurf1 and Smurf2 are closely related HECT E3s, which are 80% identical to each other. We prepared two of the most potent compounds **1** and **2** and evaluated their *in vitro* activity against Smurf1 (Figure 1A). First, we evaluated the activity of compounds **1-2** against the full length Smurf1 using our previously reported UbFluor assay.[12] UbFluor is an established robust assay that has been validated and is commonly used to quantify the enzymatic activity of HECT and RBR E3 ligases in biochemical and high throughput screening assays.[12], [13] Compound **1** and compound **2** have IC_50_’s of 230nM and 15µM (Figure 1B) respectively and compound **1** inhibition effect was further confirmed by an auto-ubiquitination assay (Supplementary Figure 1A). Notably, these IC_50_ values are significantly different from the single digit nanomolar IC_50_’s reported in the original patent. Because, UbFluor is an E2 independent activity assay, we conclude that compounds **1-2** are E2 independent Smurf1 inhibitors. We next asked the question of how these molecules might inhibit the enzyme. Smurf1 contains four domains: C2 domain, two WW domains, and HECT domain. The function of C2 domain is to bind Ca^2+^ and lipid membranes, WW domain recruits substrates to Smurf1 by recognizing PPxY motifs, while HECT domain performs the catalytic function that orchestrates poly-ubiquitination of substrates.[14] Because compounds **1-2** inhibited the catalytic activity of Smurf1 and because the catalytic HECT domain of Smurf1 is responsible for its catalytic activity, we hypothesized that compounds **1-2** might inhibit the catalytic HECT domain of Smurf1. To test this hypothesis, we prepared the isolated catalytic Smurf1 HECT domain and tested the activity of our compounds against this construct. As previously, both compounds inhibited the activity of Smurf1 in both UbFluor (compound **1** IC_50_ = 450nM and compound **2** IC_50_ **=** 1.7µM) and auto-ubiquitination assays (Figure 1C). Because autoubiquitination assays are not robust we used them for qualitative assessment of inhibitor activity, while UbFluor was used to quantify the inhibitor activity. Our experiments confirmed the original hypothesis that both compounds act by inhibiting the catalytic HECT domain of Smurf1. Interestingly, in auto-ubiquitination inhibition assays we observed an accumulation of E2∼Ub thioester, which is consistent with the inhibition of the transthiolation reaction between the E2∼Ub thioester and Smurf1 (Figure 1D). Additionally, we observed that the inhibition of Smurf1 HECT was time independent, indicating that the molecules were non-covalent inhibitors (Supplementary Figure 1B). [15]

**Figure 1.**
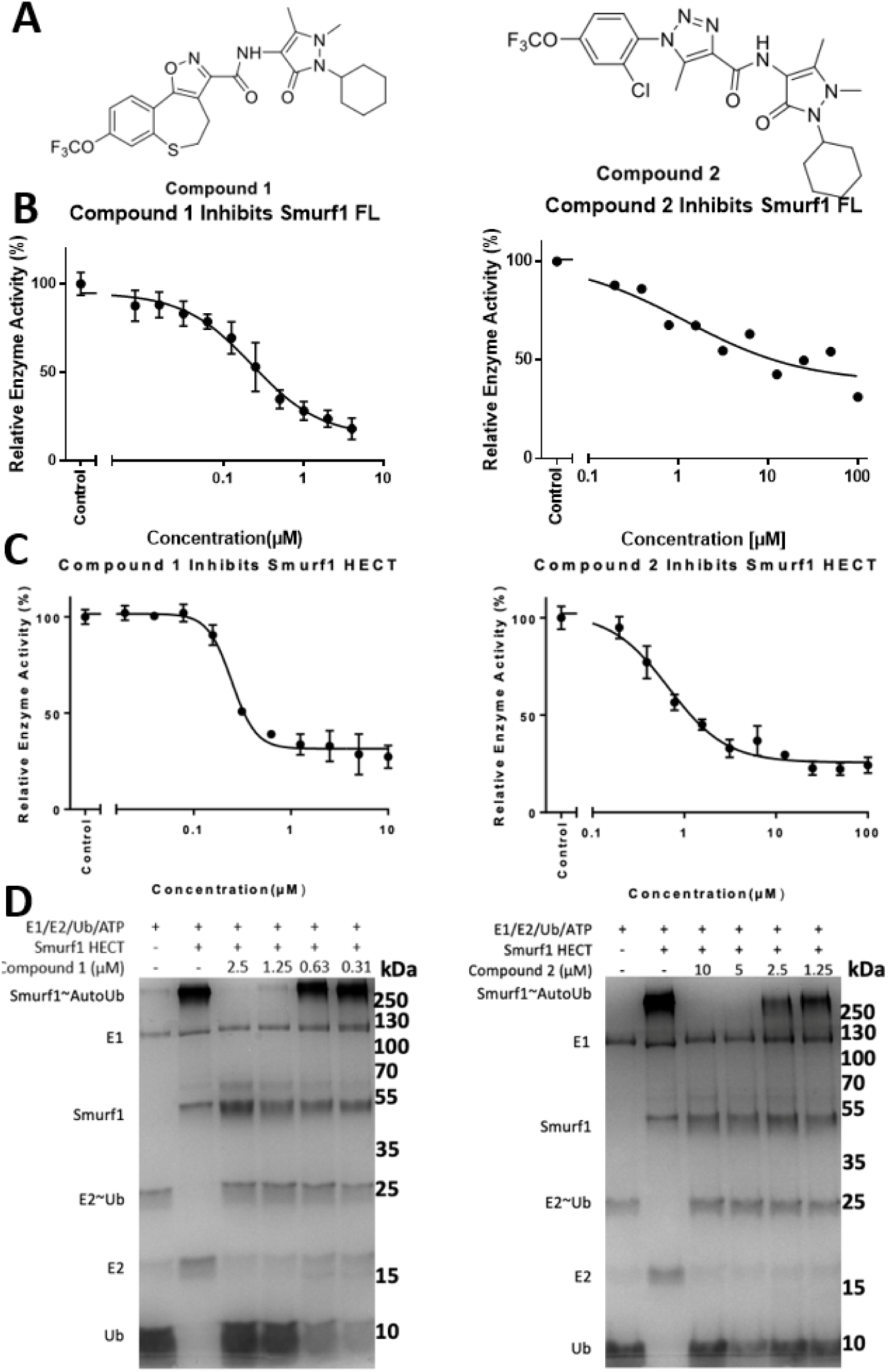
Compounds **1** and **2** inhibit Smurf1 HECT domain. **(A)** Chemical structure of compound **1** and **2**. **(B)** Compound **1** and **2** inhibits Smurf1 FL activity in UbFluor assay. Compound **1** and **2** (10µM) was serial diluted and reacted with Smurf1 FL (0.5µM) and UbFluor (2µM) and monitored via plate reader and found to have IC_50_ = 230nM and 15µM respectively. **(C)** Smurf1 HECT (0.5µM) and UbFluor (2µM) were treated with the different concentrations of compound **1** and **2**, followed by measurement of the fluorescence polarization. Plotting slopes of the linear region vs compound concentration afforded dose response curves. Compound **1** IC_50_ 450nM, compound **2** IC_50_ 1.7µM. **(D)** Compound **1** and **2** inhibits autoubiquitination of Smurf1 HECT in a dose dependent manner. His-Tag UBE1 (0.2µM), UbcH7 (2µM), Ubiquitin (10µM) and ATP (2µM) were incubated with Smurf1 HECT (2µM) in the presence or absence of compound **1** for 45 min, followed by SDS PAGE and coomassie staining.

To further confirm the possible mode of inhibition, we prepared C-terminally truncated Smurf1 HECT domain (Smurf1^trunc^) that lacks the C-terminal hydrophobic –FAVE-COOH peptide that is highly conserved among HECT family. HECT E3s that lack this C-terminal fragment can undergo a transthiolation reaction with E2∼Ub thioesters but cannot transfer ubiquitin onto the incoming lysine of the protein substrate.[16] Recent structural studies suggest that the C-terminal hydrophobic tail of HECT E3s is anchored at the N-lobe of the HECT domain, thereby stabilizing the L-conformation, which is needed for the isopeptide bond ligation step.[17] Moreover, the C-terminus of HECT E3s and the N-lobe of the HECT domain contribute to the reactivity toward the covalent activity-based ubiquitin probes, suggesting that such probes may preferentially react with the L-conformation of HECT E3 ligases.[17] Both compounds **1** and **2** inhibited the formation of Smurf1^trunc^∼Ub thioester in our assays (Figure 2A and 2B). Furthermore, the catalytic cysteine of Smurf1^trunc^ was not labeled by the C-terminal Ub propargyl (UbPA) activity-based probes, confirming that the C-terminal –FAVE peptide is essential for the covalent labeling of cysteine of Smurf1 (Figure 2C). Finally, both compounds **1** and **2** inhibited the covalent labeling of Smurf1 HECT domain by the UbPA probe, suggesting that compounds **1-2** may block the function of the –FAVE peptide (Figure 2D) and destabilize the L-conformation of the Smurf1 HECT domain. Altogether our biochemical data show that Smurf1 inhibitor mechanism of action is complex. On one end compound **1** and **2** prevent the formation of HECT Smurf1∼Ub thioester and this could be due to several mechanisms. The direct blocking of E2∼Ub/HECT Smurf1 interaction, blocking the catalytic cysteine of Smurf1, and allosteric mechanism that destabilizes the T-conformation of the HECT domain, which is required for transthiolation. On the other hand, compounds **1** and **2** prevent the covalent labeling of Smurf1 HECT domain with UbPA probe, suggesting that they can either block the catalytic cysteine or may also destabilize the L-conformation of the Smurf1 HECT domain required for the covalent probe labeling and isopeptide ligation. According to this complex mechanism, we observe the accumulation of both E2∼Ub and to a less degree Smurf1∼Ub thioesters in the native ubiquitination reaction (Figure 1D). Finally, we measured the binding of compounds **1** and **2** toward the Smurf1 catalytic HECT domain using biolayer interferometry methods and discovered that compounds **1** and **2** bind to Smurf1 HECT domain with K_d_ = 25.8µM ± 2.2µM, k_a_ = 2.7 x 10^4^ M^-1^s^-1^, k_dis_ = 6.9 x 10^-1^ s^-1^, and K_d_ = 54.5µM ± 2.0µM respectively, k_a_ = 2.1 x 10^3^ M^-1^s^-1^, and k_dis_ = 1.2. x 10^-1^ s^-1^ (Supplementary Figure 2A and 2B).

**Figure 2.**
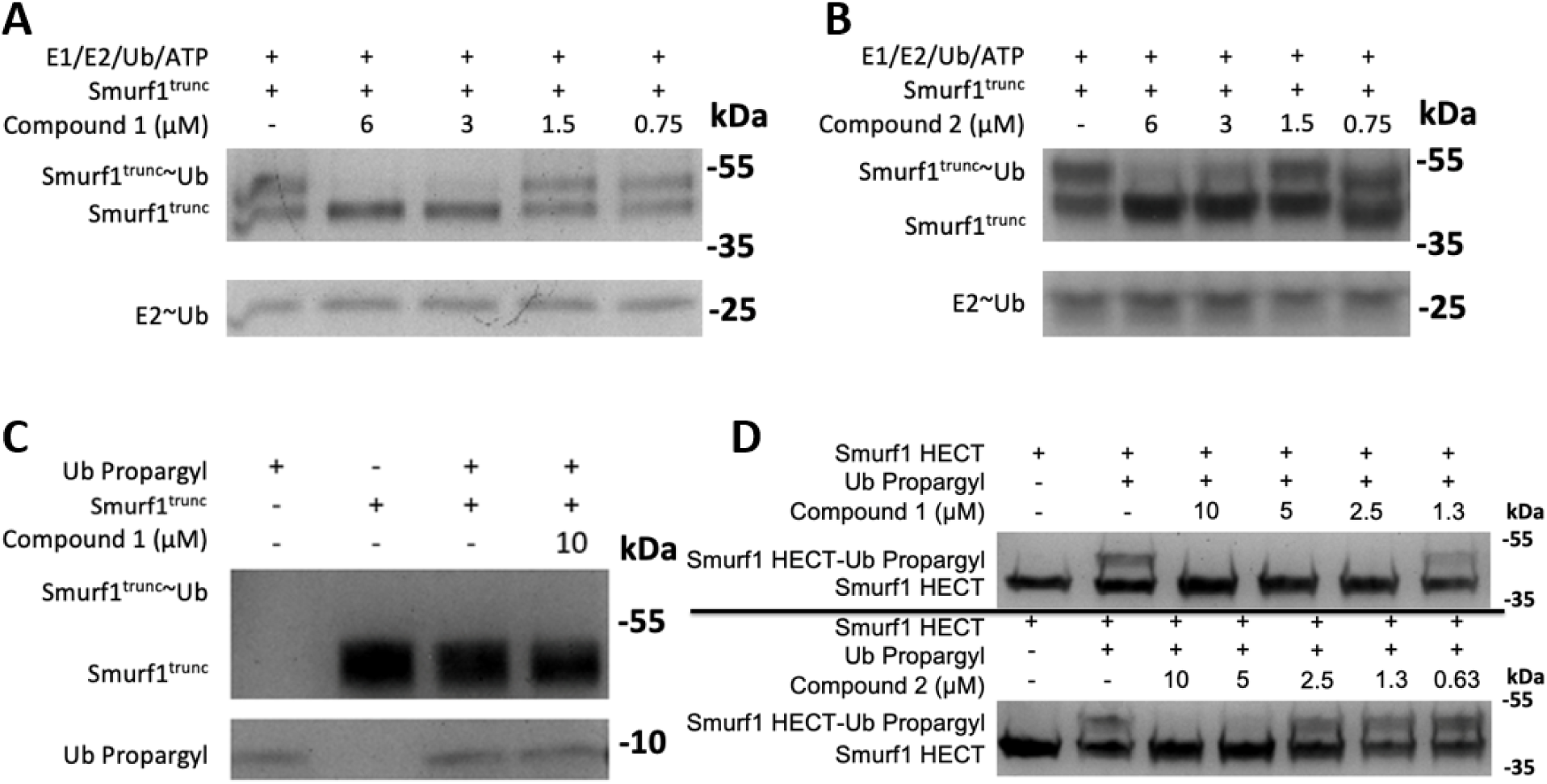
Compounds **1** and **2** inhibit E2∼Ub/Smurf1 transthiolation and Ub propargyl labeling of Smurf1. **(A)** Compound **1** prevents E2∼Ub/Smurf1^Trunc^ transthiolation. His-Tag UBE1 (0.2µM), UbcH7 (2µM), Ubiquitin (10µM) and ATP (2µM) were incubated with Smurf1^Trunc^ (2µM) in the presence and absence of compound **1** for 45 min, followed by SDS page and coomassie staining. **(B)** Compound **2** prevents E2∼Ub**/**Smurf1^Trunc^ transthiolation. Reactions we conducted as in (A). **(C)** C-terminus of Smurf1 is important for the covalent labeling of the catalytic cysteine with the activity-based probe Ub-propargyl. Smurf1^Trunc^ (2µM) was not labeled by Ub Propargyl (4µM) after 48 hrs. **(D)** Compounds **1** and **2** inhibit the labeling of Smurf1 HECT by Ub Propargyl. Smurf1 HECT (2µM) was incubated with Ub Propargyl (2.5µM) in the absence and presence of compound **1** or **2** for 24 hours and the formation of Smurf1 HECT-Ub Propargyl was monitored by SDS-PAGE and coomassie staining.

Next, we surveyed the selectivity of the discovered inhibitors **1** and **2**. Initially we decided to focus on several representative E3 ligases – Smurf2, WWP1, Nedd4-1, and NleL. Smurf2, WWP1, and Nedd4-1 are mammalian HECT E3s, while NleL is a bacterial HECT-like E3 with the catalytic cysteine. The HECT domain of Smurf2 is highly identical to Smurf1 (80%), while HECT domains of WWP1, Nedd4-1, and NleL are 53.3%, 54.1%, and 18.6%% identical to Smurf1 respectively. Accordingly, we expected that compounds **1** and **2** would inhibit Smurf2 but not Nedd4-1, WWP1 E3s, and NleL. As another control experiment, we tested our compounds against the catalytic domain of the deubiquitinating enzyme USP8. We have found that compounds **1** and **2** did not inhibit Nedd4-1, WWP1, NleL and USP8 as expected (Supplementary Figure 3A-3C), and they also did not inhibit WWP1 labeling by the Ub propargyl probe (Supplementary Figure 4). Interestingly compounds **1** and **2** also did not inhibit Smurf2 which is highly unusual, since the catalytic HECT domain of Smurf2 is highly identical to Smurf1, differing only in 17 aminoacids. It is likely that one or several aminoacids are responsible for the observed high selectivity of Smurf1 inhibitors.

The catalytic domain of HECT E3 ligases is highly conserved from *S.Cerevisiae* to humans. It consists of the C-lobe and the N-lobe linked via the flexible linker.[18] The catalytic cysteine is positioned at the C-lobe which contacts with the ubiquitin of the E2∼Ub thioester, while the N-lobe contacts the E2 enzyme of the E2∼Ub thioester.[19] From the seventeen aminoacids that are significantly different between Smurf1 and Smurf2, eight aminoacids are positioned on the C-lobe of Smurf1/2, while nine aminoacids are positioned on the N-lobe of Smurf1/2. We therefore decided to prepare a chimeric Smurf1 HECT (Smurf1C2) domain which contains Smurf2 C-lobe and Smurf1 N-lobe (Supplementary Figure 5). In doing so we hoped to obtain a quick estimate on what aminoacids might govern the observed selectivity of Smurf1 inhibitors. Interestingly, Smurf1C2 enzyme was active in UbFluor and autoubiquitination assays and was inhibited by both compounds **1** and **2** with IC_50_s similar to those of Smurf1 HECT domain (Figure 3). These results conclude that the C-lobe might not be responsible for the observed selectivity of Smurf1 inhibitors.

**Figure 3.**
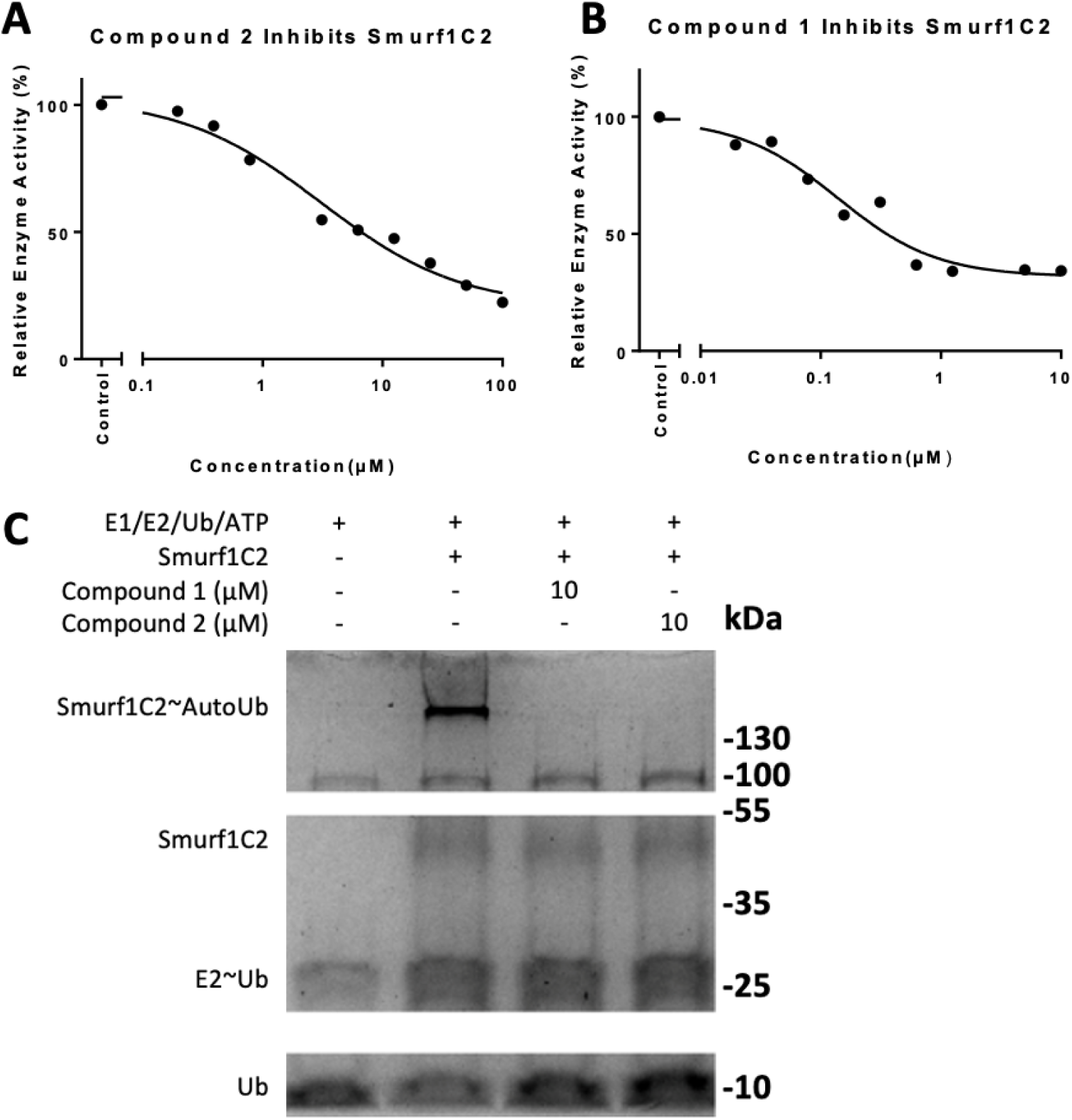
Smurf1 C-lobe is not required for selectivity against other HECT and HECT-like E3s. **(A)** Compound **1** inhibits chimera protein consisting of Smurf1 N-lobe and Smurf2 C-lobe in UbFluor assay. Smurf1C2 (0.5µM) and UbFluor (2µM) were treated with different concentration of compound **1**, followed by fluorescence polarization assay (IC_50_ 140nM). **(B)** Compound **2** inhibits Smurf1C2 activity in UbFluor assay (IC_50_ = 3.1 µM). Measurements were conducted as in (**A**). **(C)** Compound **1** and **2** inhibits autoubiquitination of Smurf1C2. His-Tag UBE1 (0.2µM), UbCH7 (2µM), Ubiquitin (10µM) and ATP (2µM) were incubated with Smurf1C2 (2µM) in the presence and absence of 10 µM compound **1** and **2** for 45 min, followed by SDS-PAGE and coomassie staining.

In summary, we report that compounds **1** and **2** are highly selective inhibitors of Smurf1, and they act by preventing the transthiolation between Smurf1 HECT and E2∼Ub thioesters. Experiments with UbPA activity-based probes showed that compounds **1** and **2** also block the covalent labeling of Smurf1 with the UbPA suggesting that they inhibit the function of the C-terminal FAVE peptide and may destabilize the L-conformation of Smurf1 HECT domain, or block the catalytic cysteine. Finally, we showed that the C-lobe of Smurf1 does not contribute to the observed specificity of Smurf1 inhibitors. The large difference between the observed K_d_ values (µM range) and potent IC_50_ values (nM range) of compound **1** and **2** is similar to that of the previously observed E2 enzyme inhibitors and requires further investigation.[20] For example, both compounds may act as glues and require Ub for a complete binding to Smurf1 HECT domain.[20] Another reason for such discrepancy is that our compounds are acting as possible aggregators in biochemical assays and selectively inactivate/destabilize Smurf1. However, we used detergents in our assays, and observed time independent IC_50_s.[21] Other factors that may rule out compound **1** and **2** as aggregators are the observed selectivity for Smurf1 in biochemical assays, and the recently reported inhibition of Smurf1 in cells by compound **1** and **2** and their analogues.[22] Additional biophysical, biochemical and structural studies are needed to fully deconvolute the exact biochemical mechanism of the reported Smurf1 inhibitors.

## Supporting Information

Protocols for protein expression, UbFluor assay, autoubiquitination assay, UbFluor data processing, Ub propargyl labeling, synthesis of compounds, and protein sequence.

## Accession IDs

Smurf1: Q9HCE7

Smurf2: Q9HAU4

Nedd4-1: P46934

WWP1: Q9H0M0

NleL: A0A3J1F006

USP8: P40818

## Acknowledgment

Research reported in this publication was supported by the National Institute of General Medical Sciences of the National Institute of Health under Award Number R01GM115632. The content is solely the responsibility of the authors and does not necessarily represent the official views of the National Institute of Health.

## Abbreviations

UbPA: Ubiquitin Propargyl

## Graphics

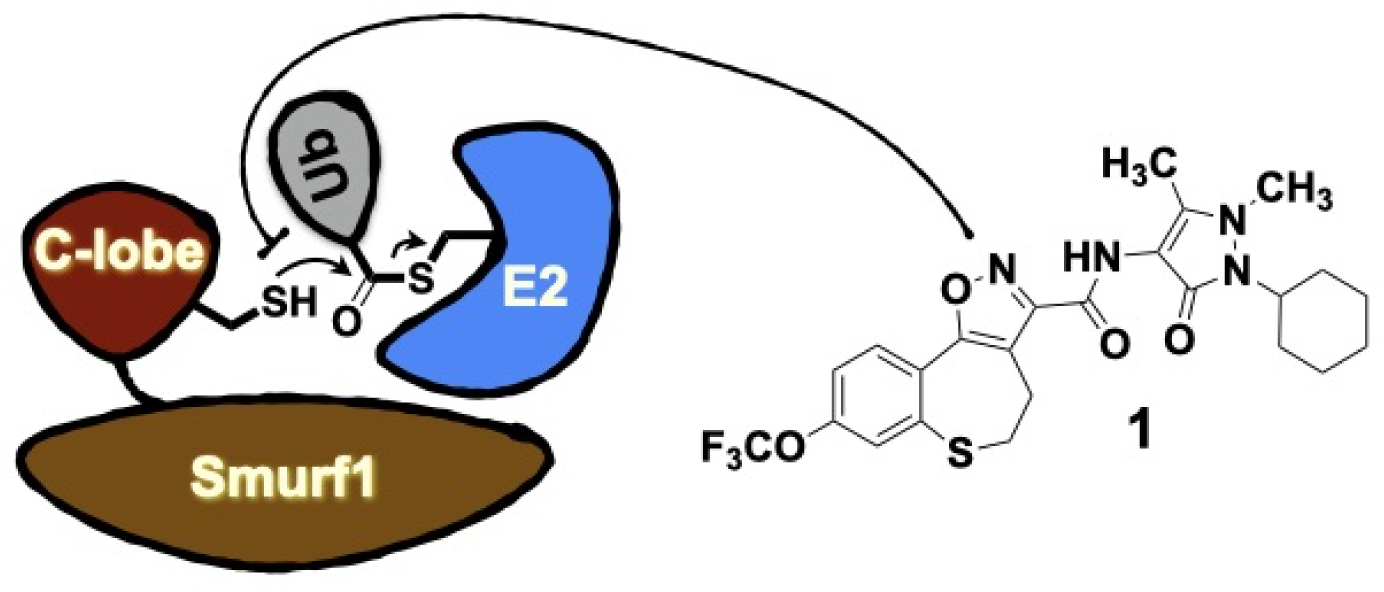

## SUPPORTING INFORMATION

### Supplementary Data

**Supplementary Figure 1.**
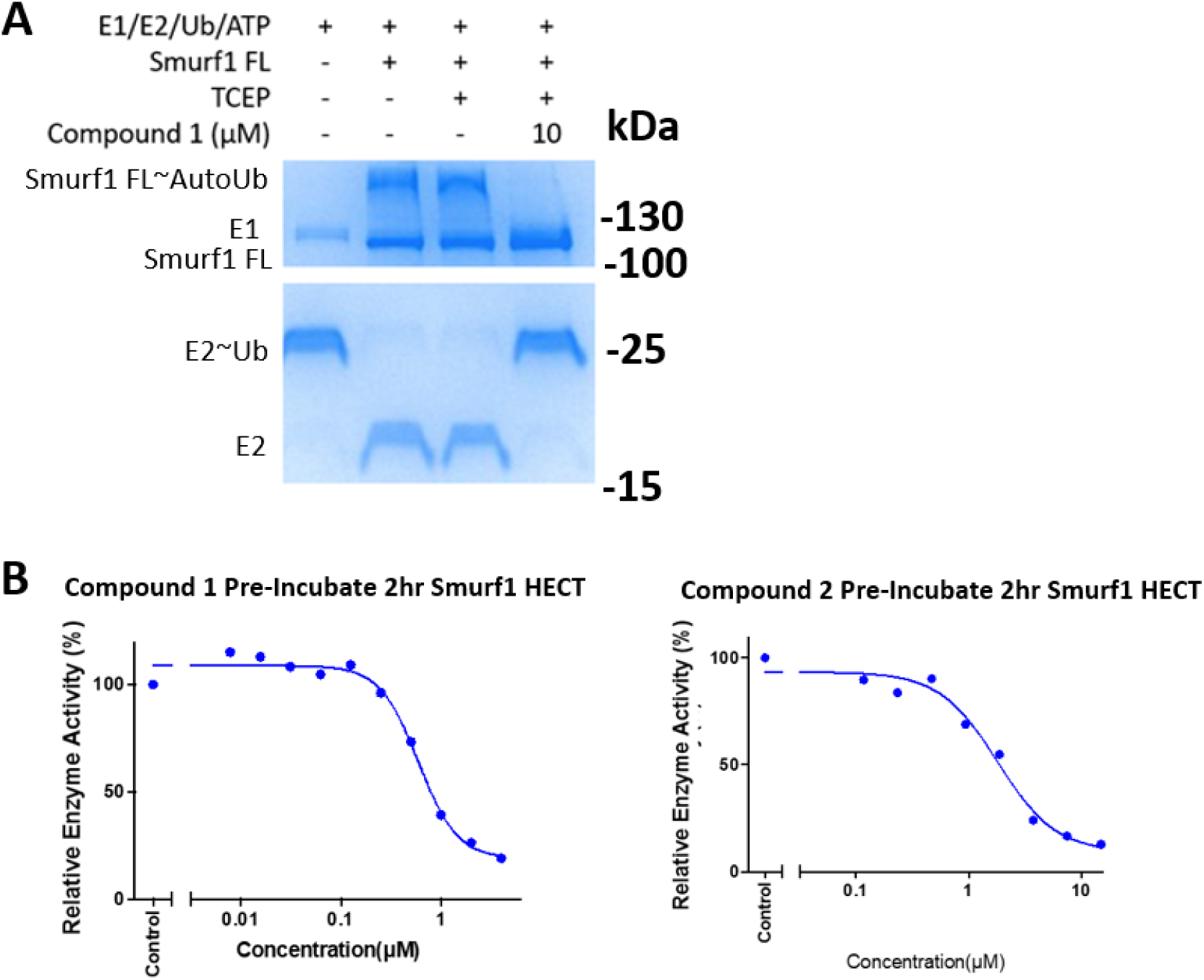
Compounds **1** and **2** inhibit full length Smurf1 FL. **(A)** Compound **1** inhibits autoubiquitination of Smurf1 FL. His-Tag UBE1 (0.2µM), UbCH7 (2µM), Ubiquitin (10µM) and ATP (2µM) were incubated with Smurf1 FL (2µM) in the presence and absence of 10 µM compound **1** for 45 min to monitor autoubiquitination. The reaction was resolved by SDS-PAGE followed by coomassie staining. (**B**) Compound **1** and **2 i**nhibits Smurf1 HECT non-covalently. Compound **1** and **2** (10µM) were serial diluted and reacted with Smurf1 HECT (0.5µM) and UbFluor (2µM) in the presence and absence of a 2hr pre-incubation time. In the presence of 2hr pre-incubation, compound **1** and **2** had IC_50_ values of 595nM and 1.8µM respectively whereas 0hr pre-incubation had very similar values of 450nM and 1.7µM.

**Supplementary Figure 2.**
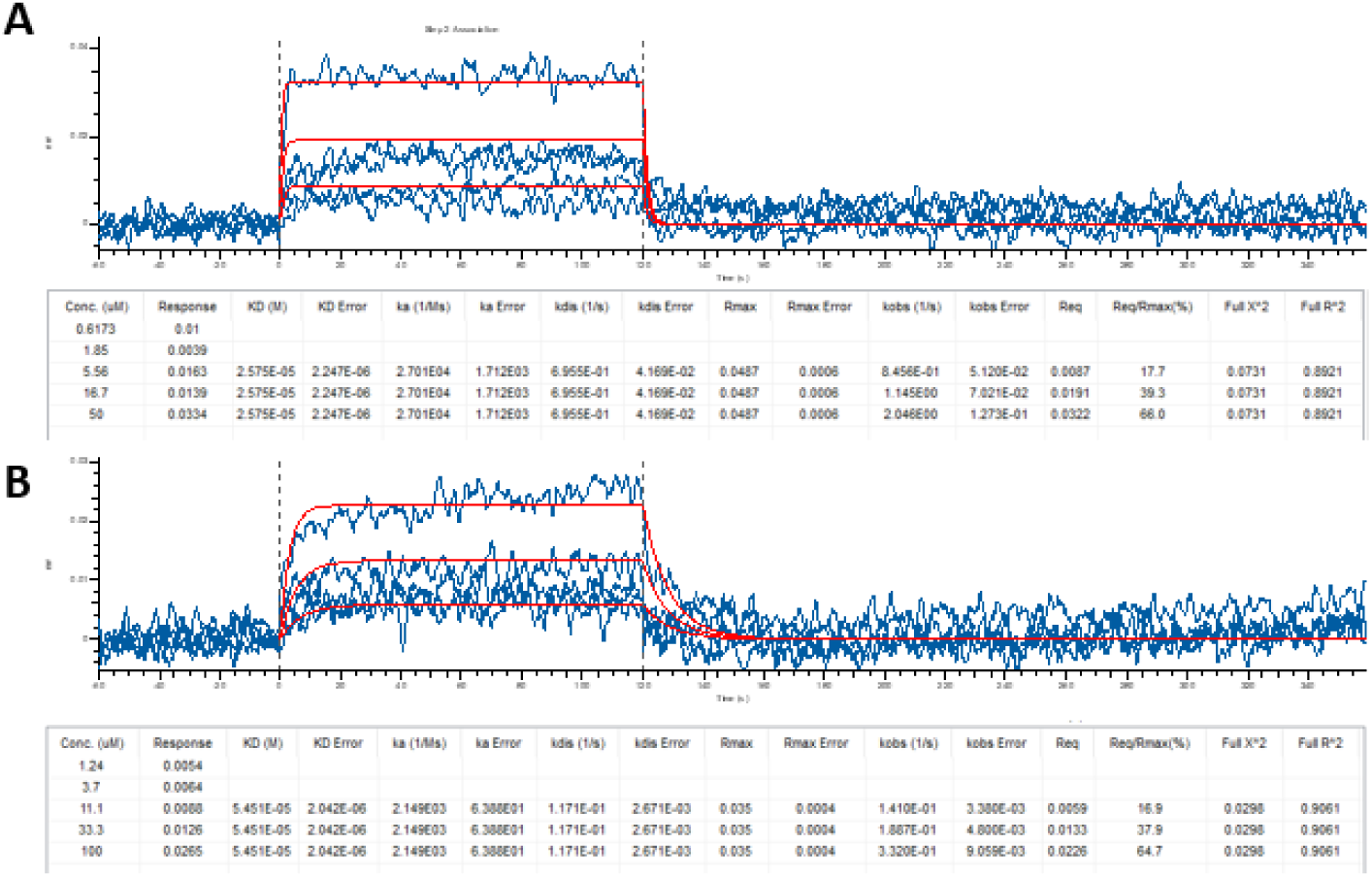
Binding of compounds **1** and **2** to Smurf1 HECT Domain as measured by Biolayer Interferometry. **(A)** Compound **1** binds to Smurf1 HECT with a K_d_ = 25.8 ± 2.2µM, K_a_ = 2.7 x 10^4^ M^-1^s^-1^, and K_dis_ = 6.9 x 10^-1^ s^-1^. Biotinylated Smurf1 HECT (40µg/mL) was immobilized onto super streptavidin biosensors and allowed to associate and dissociated with 50µM, 16.7µM, 5.6µM, 1.9µM, and 0.6µM compound **1** and K_d_ was calculated. **(B)** Compound **2** binds to Smurf1 HECT with a K_d_ = 54.5 ± 2.0µM, K_a_ = 2.1 x 10^3^ M^-1^s^-1^, and K_dis_ = 1.2. x 10^-1^ s^-1^. Biotinylated Smurf1 HECT (40µg/mL) was immobilized onto super streptavidin biosensors and allowed to associate and dissociate with 100µM, 33.3µM, 11.1µM, 3.7µM, 1.2µM Compound **2**.

**Supplementary Figure 3.**
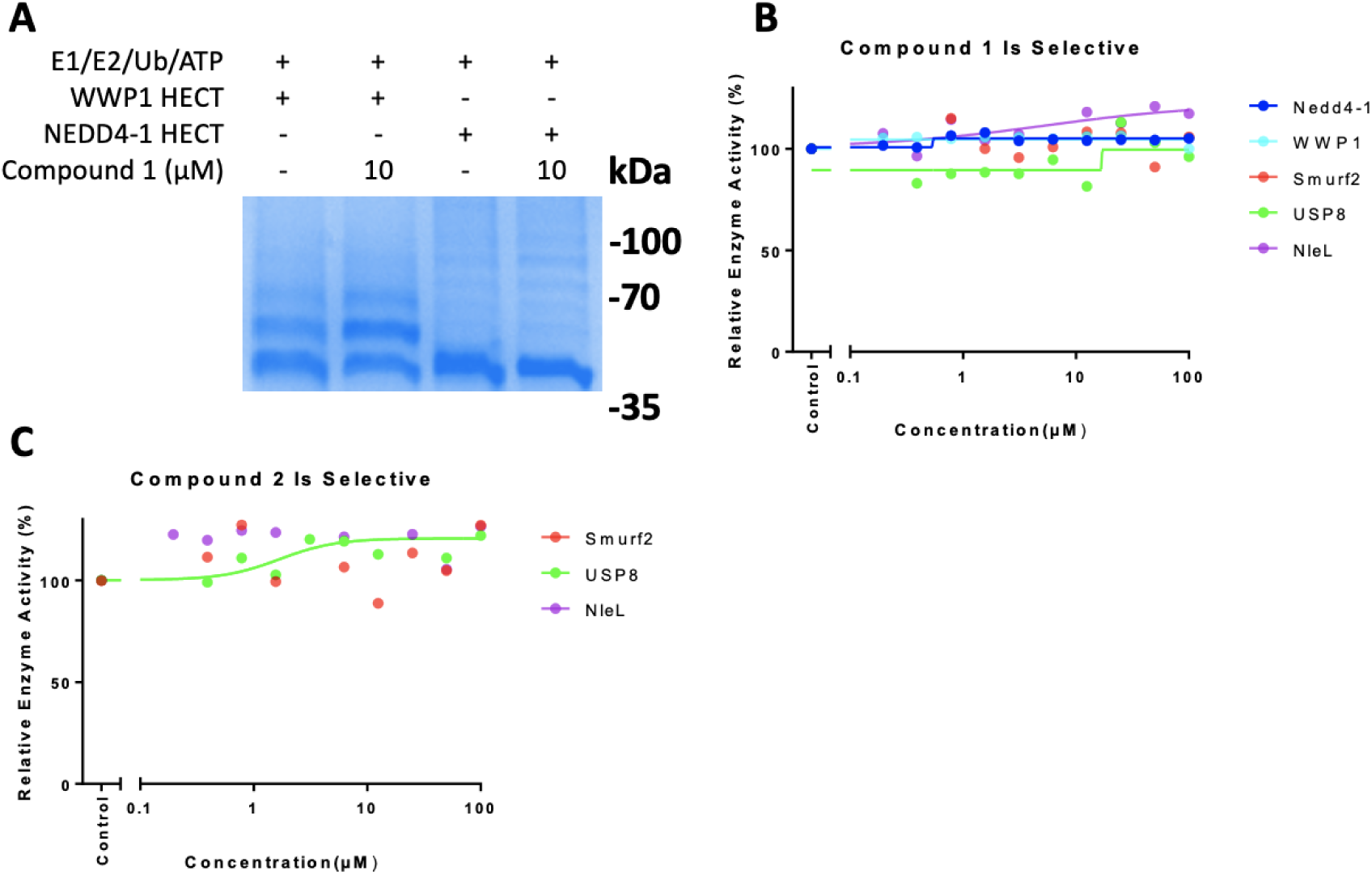
Compounds **1** and **2** do not inhibit other E3 ligases or DUBs. **(A)** Compound **1** does not inhibit autoubiquitination of WWP1 and NEDD4-1 HECT domains. His-Tag UBE1 (0.2µM), UbCH7 (2µM), Ubiquitin (10µM) and ATP (2µM) were incubated with WWP1 HECT and NEDD4-1 HECT (2µM) in the presence and absence of 10 µM Compound **1** for 45 min to monitor autoubiquitination in SDS-PAGE followed by coomassie staining. **(B)** Compound **1** does not inhibit NEDD4-1, WWP1, Smurf2, and NleL E3 ligases and USP8 DUB in UbFluor and UbTAMRA assays. **(C)** Compound **2** does not inhibit Smurf2, USP8 and NleL E3 in UbFluor and UbTAMRA assays. Experiments conducted as in (B).

**Supplementary Figure 4.**
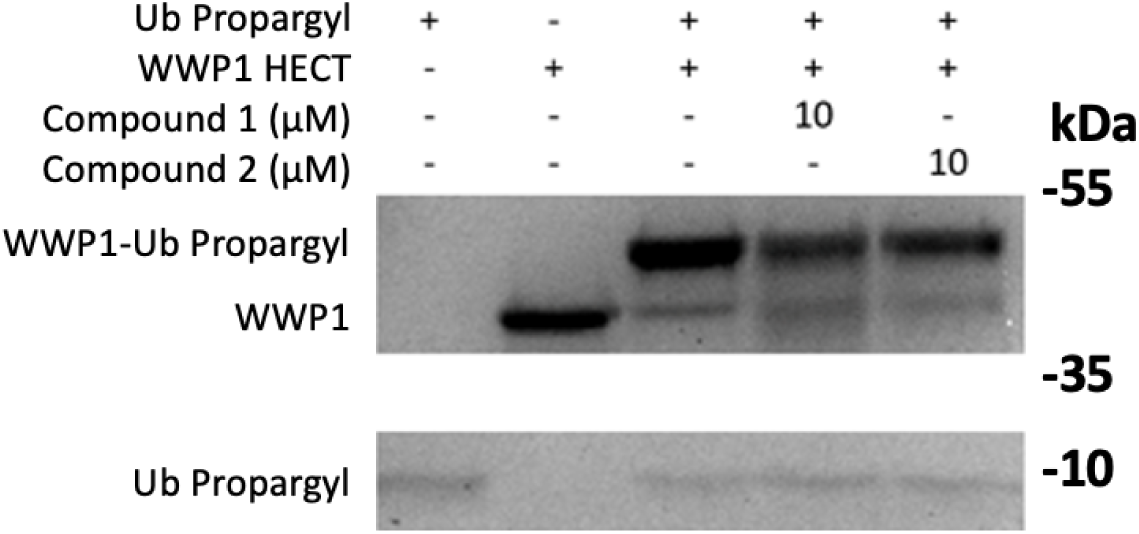
Compound **1** and **2** do no inhibit WWP1 labeling by Ub-propargyl probe. WWP1 HECT (2µM) was incubated with Ub-propargyl (2.5µM) in the absence and presence of 10 µM compound **1** and **2** for 48 hours and the formation of WWP1-Ub Propargyl was monitored by SDS-PAGE followed by coomassie staining.

**Supplementary Figure 5.**
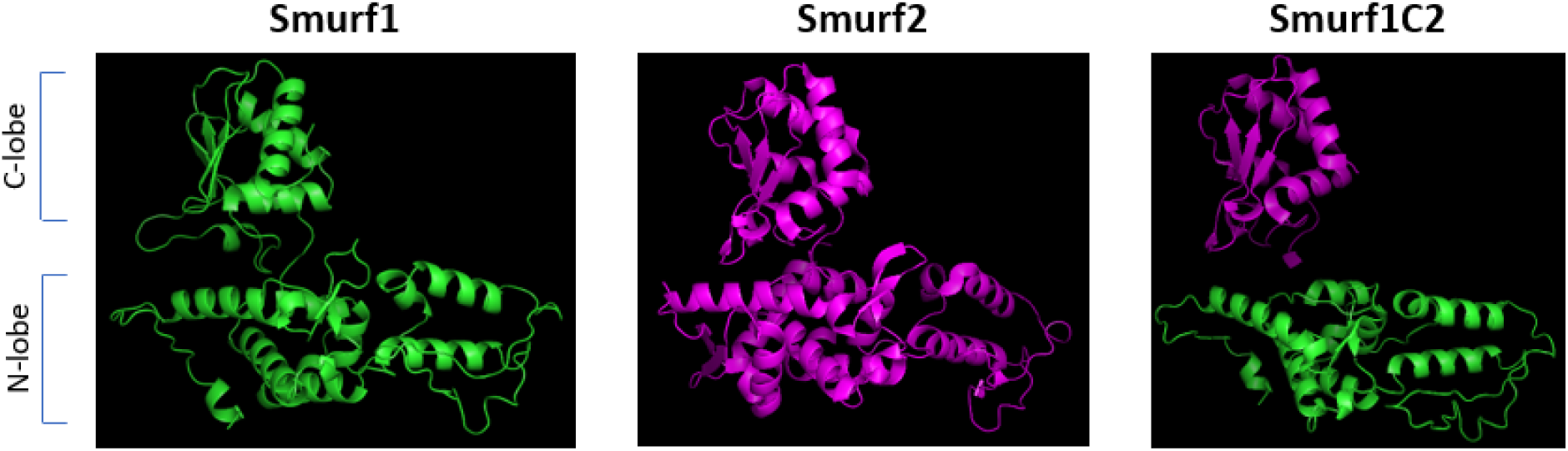
Visualization of Smurf1C2 chimera protein. Compounds **1** and **2** inhibit Smurf1 HECT but not Smurf2 HECT. The N-lobe of Smurf1 HECT was fused with the C-lobe of Smurf2 HECT to create chimeric Smurf1C2. Smurf2 HECT PDB: 1ZVD.

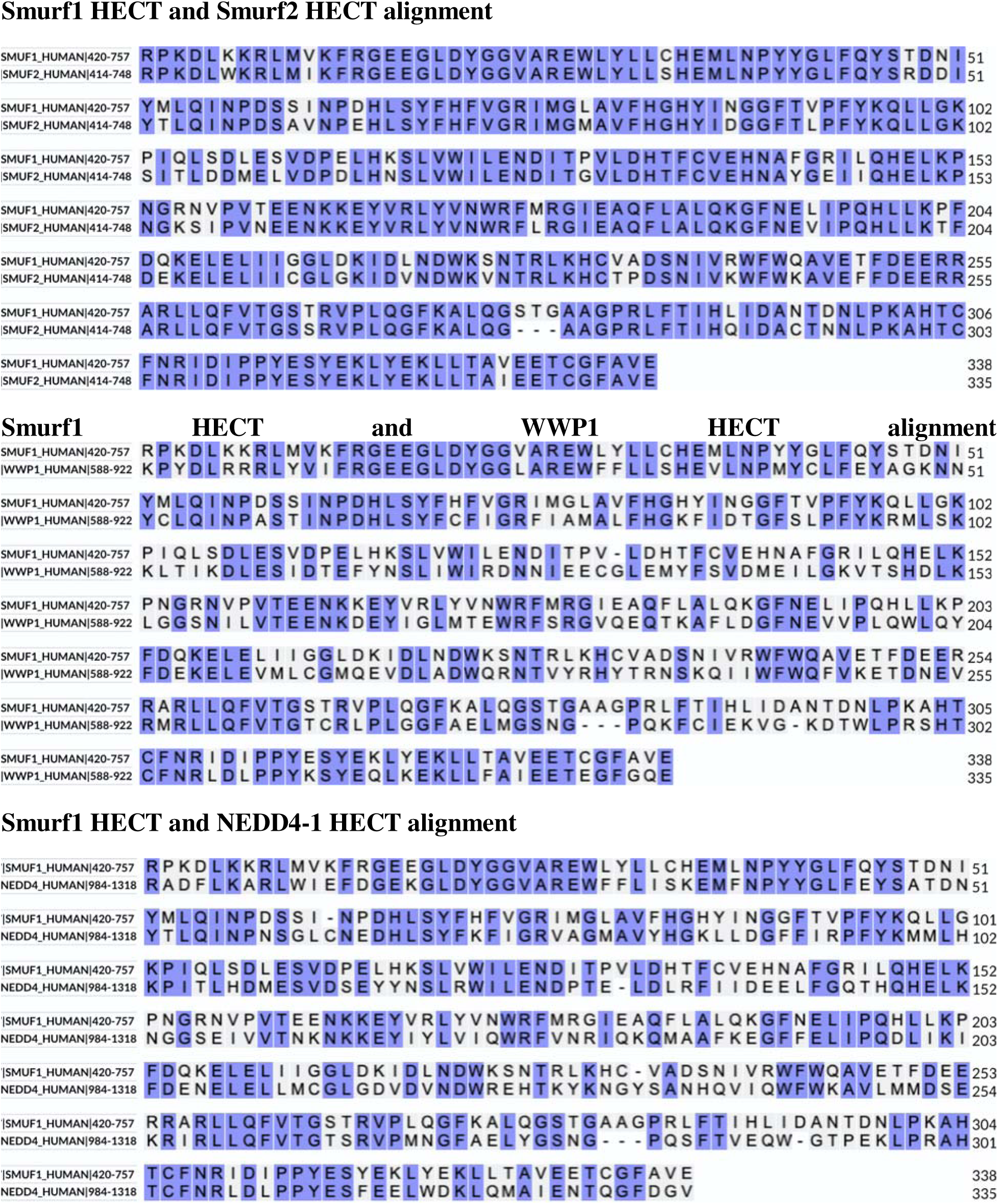

## Materials and Methods

### General Information

UBE1 (recombinant human His6), UbCH7 (recombinant human UBE2L3), NleL (E. Coli O157:H7), and Ub-Propargyl (recombinant human) were purchased from R&D systems. DNA for Smurf1, Smurf1^trunc^, Smurf1C2, and Smurf2 were synthesized by GeneArt. Gel imaging was done on a BioRad ChemiDoc MP Imaging System. All kinetic assays were done on Biotek Synergy H1 microplate reader. All Biolayer Interferometry experiments were done on Sartorius Octet R8 Protein Analysis System.

Methanol (ACS grade), ethyl acetate (ACS grade), hexane (ACS grade), dichloromethane (ACS grade), and acetonitrile (HPLC grade) were purchased from Fisher Scientific and used without further purification. Reactions were monitored by thin layer chromatography (TLC) plates (Sigma Aldrich). Flash column chromatography was performed on Teledyne Isco Combiflash RF with Redisep Rf Gold columns (Teledyne). HPLC was performed on Agilent 218 Purification System and NMR spectra was recorded on JEOL 600 MHz spectrometers.

### His-tagged Smurf1 HECT, His-tagged Smurf1^Trunc^, His-tagged Smurf1C2, His-tagged WWP1, His-Tagged Smurf2 purification

BL21 (DE3) Chemically Competent Cells (Sigma Aldrich) were transformed with His-Smurf1 in pMA-T vector and were induced with IPTG (2mM) overnight at 16°C. Cells were centrifuged and collected in phosphate buffered saline (PBS) and lysed with sonication in lysis buffer (50mM Hepes, 500mM NaCl, 1% Tween20, 5% Glycerol, half a tablet of cOmplete EDTA-free protease inhibitor cocktail [Roche]). The suspension was sonicated at 150J in intervals of 1.5secs for 20 sec total in ice. The suspension was centrifuged and the supernatant was filtered through a 20 micron PES filter.

The following purification steps were all done on the AKTA Pure (Cytiva). The resulting lysate was loaded onto a 1mL HiTrap TALON Column (Cytiva) and eluted with a gradient of 5mM imidazole buffer to 150mM imidazole buffer. The protein was then further purified by gel filtration on a HiLoad 16/600 SuperDex 75 pg column using gel filtration buffer. Elutions were quantified via SDS-PAGE electrophoresis gel. Fractions containing protein corresponding to His-Tagged Smurf1 HECT Domain protein were pooled and concentrated using Pierce Protein Concentrator PES filter. The same expression and purification steps were used for Smurf1^Trunc^, Smurf1C2, WWP1, and Smurf2.

### GST-Tagged Smurf1 Full Length, GST-tagged NEDD4-1 purification

The protein expression steps followed the same protocol as the His-Tag Smurf1 HECT Domain. The purification was done on AKTA Pure (Cytiva). The lysate was loaded onto a GSTrap 4B and eluted with a gradient of 0mM reduced glutathione buffer to 10mM reduced glutathione buffer. The protein was then further purified by gel filtration on a HiLoad 16/600 SuperDex 200 pg column using gel filtration buffer. Elutions were quantified via SDS-PAGE electrophoresis gel. Fractions containing protein corresponding to GST-Tagged Full Length Smurf1 protein were pooled and concentrated using Pierce Protein Concentrator PES filter. The same expression and purification steps were used for NEDD 4-1.

### Protocol for UbFluor and UbTAMRA assay

All UbFluor reactions were performed in buffer containing HEPES (50mM), NaCl (150mM), Tween20 (0.01%), and TCEP (1mM) pH 7.4. 100µM compounds **1** and **2** were serial diluted and 10µL was added to 10µL Smurf1 (0.5µM) and 10µL UbFluor (2µM). 20µL of the reaction was quickly transferred to a 384 well plate and centrifuged for 5 seconds at 800 rpm to remove air bubbles. Kinetic readings are taken with a Biotek Synergy H1 Microplate Reader. Readings are done at Green FP excitation 485/20 and emission 528/20 with extended gain and 7.00 mm read height. The readings were taken in intervals of 2 min for the first 20 min and then intervals of 20min for the next 8 hours. The raw polarization values are processed using Graphpad Prism software. The baseline for the background is subtracted and the slopes for each inhibitor concentration are extrapolated to give the initial velocity. The initial velocity of each inhibitor is then plotted against the inhibitor concentration to create a dose response curve. The same protocol was used for all other HECT E3s with varying enzyme concentrations. UbTAMRA (25nM) assays were used for USP8 (5nM) and were ran in a similar fashion as UbFluor assays but at an excitation of 530/25 and emission of 590/35 with a gain of 50 and 9.50 mm read height.

### Protocol for UbFluor Data Processing

Smurf1 reaction with UbFluor was calculated to behave linearly within the first two hours of reaction. The fluorescent polarization values of the first 80 minutes were plotted against time. The baseline values of the UbFluor substrate were subtracted against all wells and the slopes were calculated for each compound concentration measured. The calculated slopes were then plotted against tested concentration values and fitted against a nonlinear regression curve to get IC_50_ values. Relative Enzyme Activity is defined as the enzyme’s ability to consume UbFluor at any given concentration relative to control groups. 100% consumption of UbFluor is determined by reaction between E3 ligase and UbFluor in the absence of inhibitors and 0% consumption of UbFluor is determined by UbFluor in the absence of E3 ligase and inhibitors.

### Protocol for autoubiquitination assay for Smurf1

All autoubiquitination assays for Smurf1 were done in buffer containing HEPES (25mM), NaCl (100mM), MgCl_2_ (4mM), TCEP (1mM) pH 7.4. His-Tag UBE1 (0.2µM), UbCH7 (2µM), Smurf1 (2µM), Ubiquitin (10µM) and ATP (2µM) were added in the presence and absence of a serial dilution of compound **1** (2.5µM) and compound **2** (10µM). The reaction was incubated at room temperature for 45 min and quenched with 4x laemmli buffer. The reaction was boiled at 90°C for 10 min and loaded onto a 12% SDS-PAGE gel and electrophoresis was ran at 150V for 50 min. The gel was washed for 30 min with 100 mL water, stained for 1 hour with 25mL EZ-Blue Gel Staining Reagent, and de-stained overnight with 100 mL water all done over gentle rocking. The gel was visualized with a BioRad ChemiDoc MP Imaging System.

### Protocol for Biolayer Interferometry

All biolayer interferometry experiments were ran on the Octet R8 Protein Analysis System. Smurf1 HECT NT protein was biotinylated using the No-Weight NHS-PEO_4_-Biotin kit (Thermo Fisher) due to its compatibility with Super Streptavidin (SSA) biosensors. The SSA sensors were first equilibrated in BLI buffer (20 mM Hepes, 100 mM NaCl, and 0.1% Tween20) for 10 minutes. The sensors were equilibrated again in BLI buffer for 5 min to establish a baseline. The sensors were then dipped into the loading well containing 50µg/mL Biotin-Smurf1 for 10 min until saturation of binding. The SSA sensors were dipped into Biocytin (20 µg/mL) for 1 min until all unbound receptors on SSA sensors were blocked off. The SSA sensors were dipped into a washing well followed by an equilibration well to dissociate any remaining unbound Biotin-Smurf1 and Biocytin from the sensor for 4 min and 2 min respectively. The sensors were dipped into DMSO buffer well (20mM Hepes, 100mM NaCl, 0.1% Tween20, and 1% DMSO) for 1 min to establish the baseline for association. The sensors were dipped into the initial analyte well (0.617µM Compound 1) for 2 min and association was measured. The sensors were dipped back into DMSO buffer well for 4 min and dissociation was measured. The process of dipping the SSA sensors into baseline, association, then dissociation repeated for each remaining analyte well containing 1.85, 5.56, 16.7, and 50µM Compound 1. Compound 2 was ran with the same protocol using 1.24, 3.7, 11.1, 33.3, and 100 µM Compound 2 instead.

### Protocol for Ub-PA labeling

All Ub-PA labeling experiments were done in buffer containing HEPES (25mM), NaCl (100mM), MgCl_2_ (4mM), TCEP (1mM) pH 7.4. Smurf1 (2µM) and UbPA (4µM) was incubated for 24 hours in the presence and absence of 2x serial dilution of compound **1** and **2** (10µM). The reaction was quenched with 4x laemmli buffer and boiled at 90°C for 10 min. The reaction was loaded onto a 12% SDS-PAGE gel and electrophoresis was ran at 150V for 50 min. The gel was washed for 30 min with 100 mL water, stained for 1 hour with 25mL EZ-Blue Gel Staining Reagent, and de-stained overnight with 100 mL water all done over gentle rocking. The gel was visualized with a BioRad ChemiDoc MP Imaging System.

### Synthesis of Compound 1

The following synthesis was done following the synthetic steps found in Patent Publication No. US2017/0088546 A1.

### Compound 1 Synthesis Scheme

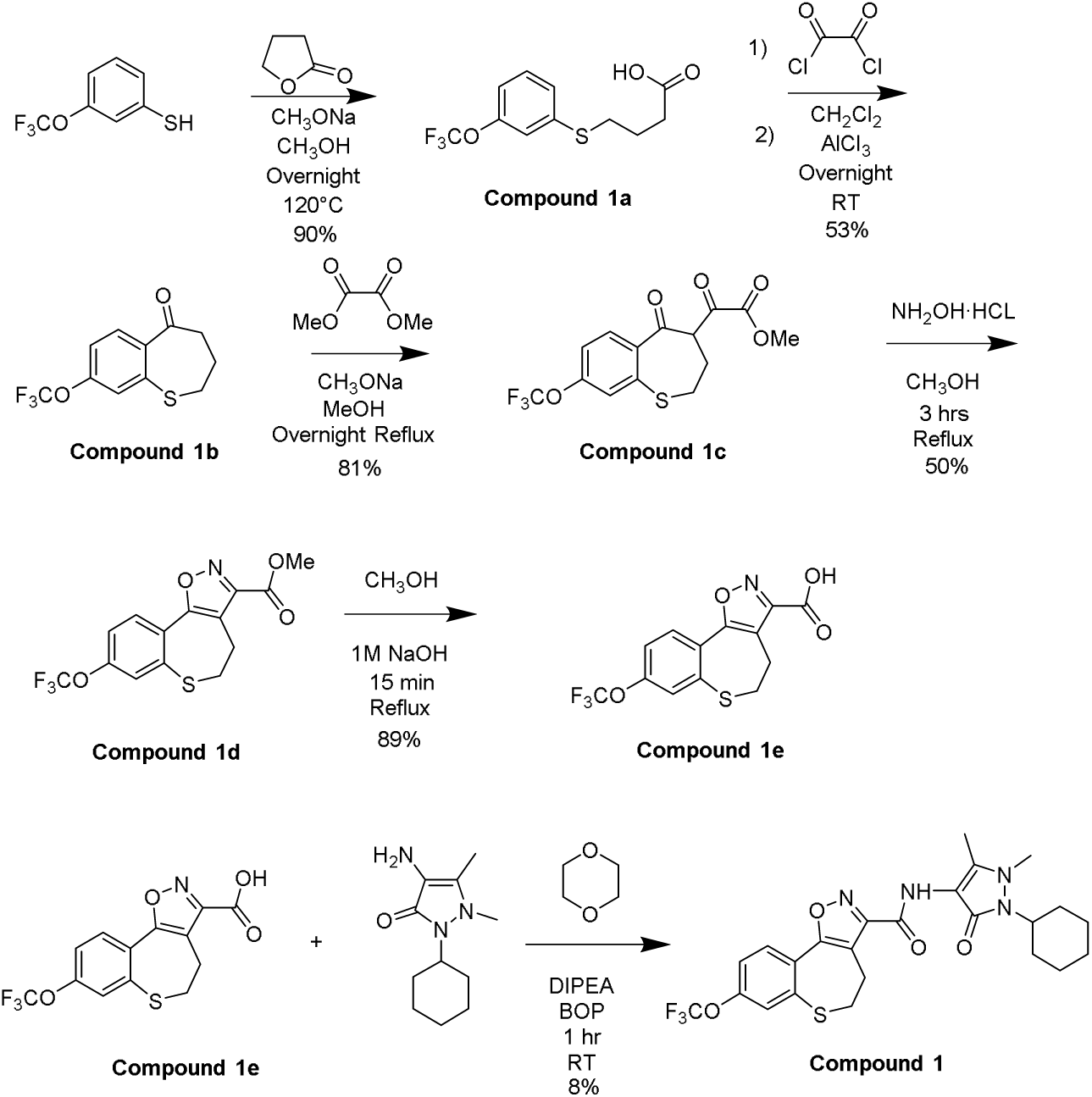

### Synthesis of Compound 1a

Sodium methoxide (278mg, 5.2mmol) was added to a 100mL round bottom flask under nitrogen. The flask was then equipped with a stir bar and 13 mL methanol (MeOH) was added and allowed to stir at 200rpm. 3-(Trifluoromethoxy)thiophenol (1g, 5.2mmol) was then added to the reaction vial. γ-butyrolactone (440mg, 390 mL, 5.2mmol) was added and the mixture was heated at reflux for 2 hours under nitrogen gas. The methanol was then removed via rotovap and the reaction was ran for 22 hours overnight at 120°C under vacuum. The mixture was quenched with water and acidified by dropwise addition of 12M HCl to pH = 1. The reaction was extracted with 20mL ethyl acetate (EtOAc) three times and all the organic layers were combined. The organic layer was dried with magnesium sulfate and filtered through a borosilicate filter funnel. The organic layer was concentrated via roto-evaporator and the liquid was loaded into a 12g RediSep Gold Silica Gold Disposable Column on the Teledyne Isco Combiflash system. The column was ran on a gradient of 0 to 60% EtOAc:Hexane and the product eluted around 15% EtOac to provide 1.3g of compound **1a** (90% yield). ^1^H NMR (600 MHz, CDCL_3_) δ7.30 (1H, d) 7.25-7.18 (2H, m), 6.99 (1H, ddd), 2.97 (2H, t), 2.40 (2H,d), 1.87(2H, tt). NMR matched previously reported compound.

### Synthesis of Compound 1b

Compound **1a** (1.3g, 4.6mmol) was added to a 100mL round bottom flask with a magnetic stir bar and cooled to 0°C. Oxalyl chloride (2.26mL, 23mmol) was added and the reaction mixture stirred for 2 hours at room temperature (RT) at 200rpm under nitrogen. The mixture was then evaporated via roto-evaporator and dissolved in 40mL dry dichloromethane (DCM). Aluminum chloride (1.38g, 10mmol) was added to a 250mL round bottom flask and put into a suspension with 90mL dry DCM. Previous mixture with compound **1a** was then added dropwise to the aluminum chloride suspension over 4 hours and allowed to run overnight for 21 hours at RT under nitrogen at 300rpm. The reaction was quenched with water and extracted with 30mL DCM three times and all the organic layers were combined. The organic layer was dried with magnesium sulfate and filtered through a borosilicate filter funnel. The organic layer was concentrated via roto-evaporator and liquid loaded into a 12g RediSep Gold Silica Gold Disposable Column on the Teledyne Isco Combiflash system. The column was ran on a gradient of 0 to 40% EtOAc:Hexane and the product eluted around 10% EtOAc to give 0.65g of compound **1b** (53%). ^1^H NMR (600 MHz, CD_3_OD) δ7.21 (1H, d), 7.02(1H, s) 6.82 (1H, dd) 3.10 (2H, t) 3.00 (2H, t) 2.25(2H, m). NMR matched previously reported compound.

### Synthesis of Compound 1c

NaOMe (300mg, 5.6mmol) was added to a 25 mL round bottom flask with a magnetic stir bar and dissolved with 1mL MeOH. Dimethyl oxalate (0.6g, 5.1mmol) was added to the flask to create a suspension. Compound **1b** (0.65g, 2.47mmol) was dissolved in 2.5mL MeOH and added to the suspension and the reaction was stirred for 2hrs at RT under nitrogen at 320 rpm. The reaction was then poured onto citric acid (2.5g, 13mmol) in 25mL ice water and extracted with 20mL ethyl acetate (EtOAc) three times and all the organic layers were combined. The organic layer was dried with magnesium sulfate and filtered through a borosilicate filter funnel. The organic layer was concentrated via roto-evaporator and liquid loaded into a 12g RediSep Gold Silica Gold Disposable Column on the Teledyne Isco Combiflash system. The column was ran on a gradient of 0 to 30% EtOAc:Hexane and the product eluted around 12% EtOAc to give 0.7g of compound **1c** (81% yield). ^1^H NMR (600 MHz, CD_3_OD) δ7.72 (1H, d) 6.86(1H, s) 6.67 (1H, dd) 3.28 (2H, m), 2.95-2.92 (4H, m) 2.17(2H, m). NMR matched previously reported compound.

### Synthesis of Compound 1d

Compound **1c** (0.7 g, 2 mmol) and hydroxyl amine hydrochloride (0.28 g, 4 mmol) were added to a 25 mL round bottom flask and equipped with a magnetic stir rod. 4 mL MeOH was added to the flask and the reaction was heated for 30 minutes at reflux under nitrogen at 200 rpm. The reaction was cooled to RT and the precipitate was then filtered and washed with MeOH. The precipitate was dry loaded into a 12 g RediSep Gold Silican Gold Disposable Column on the Teledyne Isco Combiflash system. The column was ran on a gradient of 0 to 100% DCM:Hexane and the product eluted around 100% DCM to give 0.35 g of compound **1d** (50% yield). ^1^H NMR (600 MHz, CD_3_OD) δ8.11 (1H, d) 7.58(1H, s) 7.43 (1H, dd) 3.99 (3H, s) 3.39 (2H, m) 3.12(2H, m). NMR matched previously reported compound.

### Synthesis of Compound 1e

Compound **1d** (0.35g, 1 mmol) was dissolved in 5mL MeOH and added to a 25 mL round bottom flask equipped with a magnetic stir bar. 1M NaOH (5mL, 5.00 mmol) was added slowly over 5 minutes. The reaction was stirred for 30 minutes at RT under nitrogen at 250 rpm. The reaction was then quenched with water and extracted with 25 mL EtOAc twice and the organic layer was thrown away. 12M HCl was then added dropwise to aqueous layer until pH = 1 and then extracted again with 25mL EtOAc three times and the organic layers were combined. The organic layer was dried with magnesium sulfate and filtered through a borosilicate filter funnel. The organic layer was concentrated via roto-evaporator to give 0.3 g of compound **1e** (89% yield), which was used in the next step without purification.

### Synthesis of Compound 1

Compound **1e** (0.1 g, 0.3 mmol) was dissolved in 2 mL 1,4-dioxane and added to a 25 mL round bottom flask equipped with a stir rod. 200 µL of N,N-Diisopropylethylamine, benzotriazole-1-yl-oxy-tris-(dimethylamino)-phosphonium hexafluorophosphate (400mg, 0.9mmol), and 4-amino-2-cyclohexyl-1,5-dimethyl-1H-pyrazol-3(2H)-one (0.1g, 0.48mmol) were added to the flask. The reaction was stirred for 2 hr at RT under nitrogen at 220 rpm. The mixture was quenched with water and extracted with 20mL Ethyl Acetate (EtOAc) three times and all the organic layers were combined. The organic layer was dried with magnesium sulfate and filtered through a borosilicate filter funnel. The organic layer was concentrated via roto-evaporator and dissolved in 1 mL water and 1 mL acetonitrile. The solution was processed through an Agilent HPLC system with a gradient of 0 to 100% acetonitrile: water over 30 minutes. The compound eluted around 25 min. The solvent was removed by lyophilization to give 12 mg of compound **1** (8% yield). ^1^H NMR (600 MHz, CD_3_OD) δ8.09 (1H, d), 7.42 (1H, s), 7.40 (1H, d) 4.06 (1H, tt), 3.41 (2H, m), 3.30 (3H, s), 3.21 (2H, m), 2.20 (3H, s), 2.07(2H, m), 1.77 (2H, br, d), 1.55 (3H, m), 1.40-1.20 (3H, m). MS m/z 523 [M+H]^+^. NMR and MS matched previously reported compound.

### Synthesis of Compound 2

The following synthesis was done following the synthetic steps found in Patent Publication No. US 9,403,833 B2.

### Compound 2 Synthesis Scheme

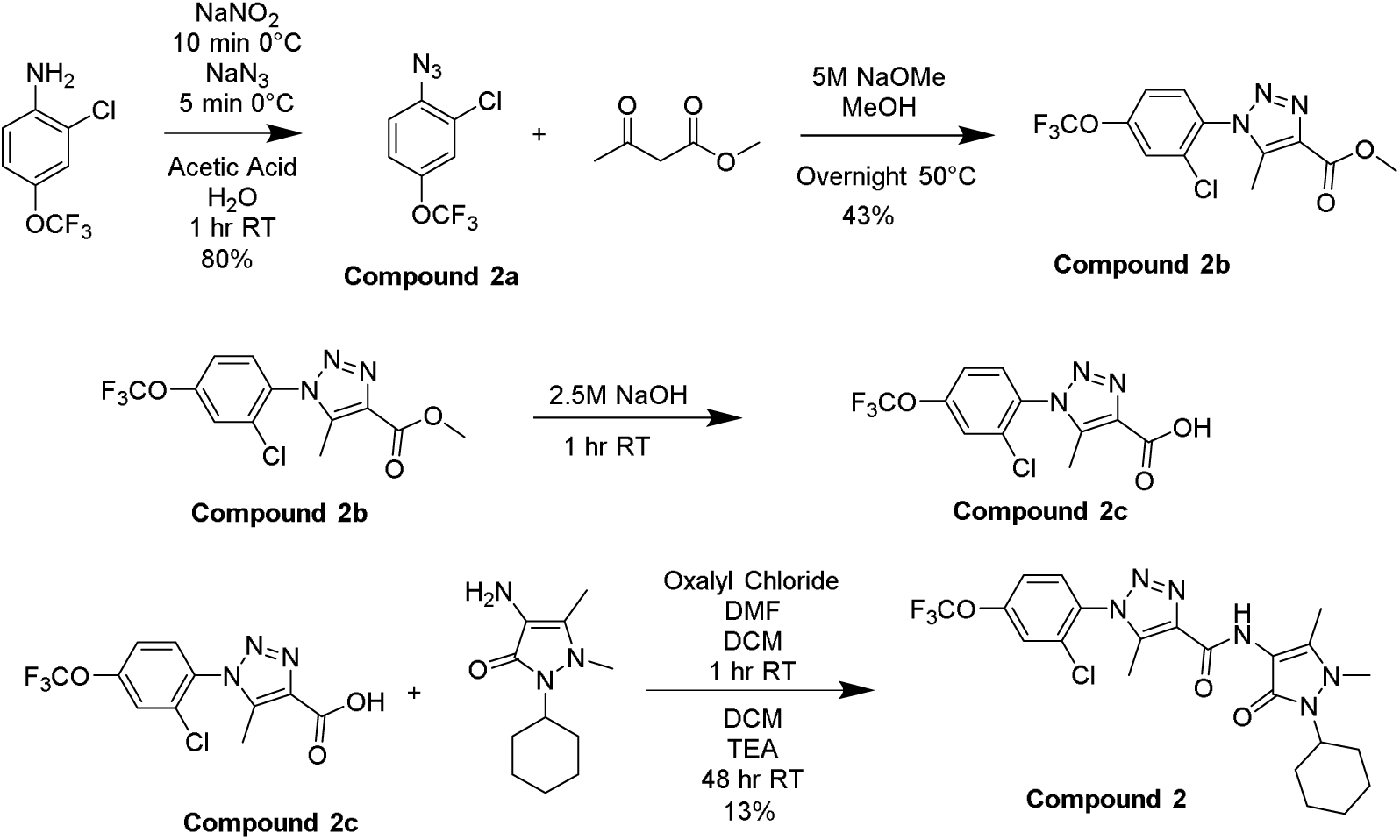

### Synthesis of Compound 2a

2-Chloro-4-(trifluoromethoxy) aniline (0.4 g, 1.9 mmol) was dissolved in 25 mL acetic acid and 25 mL water in a 250 mL round bottom flask equipped with a stir bar and cooled to 0°C in an ice bath. A solution of NaNO_2_ (0.18 g, 2.6 mmol) in 4mL water was added dropwise to the flask over 10 minutes and the reaction was stirred for an additional 10 min at 0°C at 200 rpm. A solution of NaN_3_ (0.18 g, 2.8 mmol) in 4 mL water was added dropwise over 10 minutes in the ice bath. The reaction was then removed from the ice bath and allowed to stir at RT for 30 mins at 200 rpm. The reaction was then put back on ice and then 2.5M NaOH was added dropwise until pH = 10. The reaction was extracted with 30mL ethyl acetate (EtOAc) three times and all the organic layers were combined. The organic layer was dried with magnesium sulfate and filtered through a borosilicate filter funnel. The organic layer was concentrated via roto-evaporator and liquid loaded into a 12g RediSep Gold Silica Gold Disposable Column on the Teledyne Isco Combiflash system. The column was ran on a gradient of 0 to 30% EtOAc:Hexane and the product eluted around 8% EtOAc to give 0.36g of compound **2a** (80% yield). ^1^H NMR (600 MHz, CDCl_3_) δ7.08 (1H, d), 6.90 (1H, d), 6.79 (1H, dd). NMR matched previously reported compound.

### Synthesis of Compound 2b

Compound **2a** (360 mg, 1.5 mmol) and methylacetoacetate (0.6 mL, 5.3 mmol) were dissolved in 20mL MeOH added to a 100 mL round bottle flask equipped with a magnetic stir bar. A solution of 5M NaOMe (2mL) was added to the reaction dropwise. The reaction mixture was stirred overnight for 16 hours at reflux under nitrogen at 220 rpm. The reaction was quenched with water and extracted with 30mL ethyl acetate (EtOAc) three times and all the organic layers were combined. The organic layer was dried with magnesium sulfate and filtered through a borosilicate filter funnel. The organic layer was concentrated via roto-evaporator and liquid loaded into a 12g RediSep Gold Silica Gold Disposable Column on the Teledyne Isco Combiflash system. The column was ran on a gradient of 0 to 50% EtOAc:Hexane and the product eluted around 30% EtOAc to give 0.22g of compound **2b** (43% yield). ^1^H NMR (600 MHz, CDCl_3_-d) δ7.49 (1H, d), 7.48 (1H, d), 7.32 (1H, dd), 3.95 (3H, s), 2.43 (3H, s). NMR matched previously reported compound.

### Synthesis of Compound 2

Compound **2b** (0.22 g, 0.66 mmol) was dissolved in 20mL MeOH and added to a 100mL round bottom flask equipped with a magnetic stir bar. A solution of 2.5M NaOH (2mL) was added dropwise over 10 mins and the reaction stirred for 30 min at RT at 250 rpm. The reaction was then quenched with water and extracted with 25 mL EtOAc. 12M HCl was then added dropwise to the aqueous layer until pH = 1. The aqueous layer was then extracted with 25mL EtOAc three times and the organic layers were combined. The organic layer was dried with magnesium sulfate and filtered through a borosilicate filter funnel. The organic layer was concentrated via roto-evaporator and the crude compound **2c** produced was used directly for the next step.

Crude compound **2c** (0.1 g, 0.31 mmol) was dissolved in 2mL dry DCM and added to a 10mL round bottom flask. Oxalyl chloride (0.1 mL, 1.2mmol) and dimethylformamide (0.1 mL) were added and the reaction stirred for 1 hr at RT under nitrogen at 220 rpm. 4-Amino-2-cyclohexyl-1,5-dimethyl-1H-pyrazol-3(2H)-one (0.1 g, 0.48 mmol) was dissolved in 1 mL DCM and added to the flask. Triethylamine (0.2 mL, 1.4 mmol) was then added dropwise to the flask and the reaction stirred for 48 hrs in RT under nitrogen at 220 rpm. The reaction was quenched with water and extracted with 30mL ethyl acetate (EtOAc) three times and all the organic layers were combined. The organic layer was dried with magnesium sulfate and filtered through a borosilicate filter funnel. The organic layer was concentrated via roto-evaporator and dissolved in 1 mL water and 1 mL acetonitrile. The solution was processed through an Agilent HPLC system on a gradient of 0 to 100% acetonitrile: water over 30 minutes. The compound eluted around 25 min. The solvent was removed by lyophilization. to give 20 mg product of compound **2** (13% yield). ^1^H NMR (600 MHz, DMSO-d6) δ9.32 (1H, s), 8.00 (1H, d), 7.92 (1H, d), 7.69 (1H, mult), 3.89 (1H, mult), 3.30 (3H, s), 2.33 (3H, s), 2.01 (3H, s), 2.01-1.96 (2H, mult), 1.75 (2H, mult), 1.642 (3H, mult), 1.26 (2H, mult), 1.13 (1H, mult). MS m/z 513 [M+H]^+^. NMR matched previously reported compound.

## Protein Sequences

### GST-Tag Smurf1 Full Length Protein Sequence

MSPILGYWKIKGLVQPTRLLLEYLEEKYEEHLYERDEGDKWRNKKFELGLEFPNLPYYI DGDVKLTQSMAIIRYIADKHNMLGGCPKERAEISMLEGAVLDIRYGVSRIAYSKDFETLK VDFLSKLPEMLKMFEDRLCHKTYLNGDHVTHPDFMLYDALDVVLYMDPMCLDAFPKLVCFKKRIEAIPQIDKYLKSSKYIAWPLQGWQATFGGGDHPPKSDMSNPGTRRNGSSIKIRLTVLCAKNLAKKDFFRLPDPFAKIVVDGSGQCHSTDTVKNTLDPKWNQHYDLYVGKTDSITISVWNHKKIHKKQGAGFLGCVRLLSNAISRLKDTGYQRLDLCKLNPSDTDAVRGQIVVSLQTRDRIGTGGSVVDCRGLLENEGTVYEDSGPGRPLSCFMEEPAPYTDSTGAAAGGGNCRFVESPSQDQRLQAQRLRNPDVRGSLQTPQNRPHGHQSPELPEGYEQRTTVQGQVYFLHTQTGVSTWHDPRIPSPSGTIPGGDAAFLYEFLLQGHTSEPRDLNSVNCDELGPLPPGWEVRSTVSGRIYFVDHNNRTTQFTDPRLHHIMNHQCQLKEPSQPLPLPSEGSLEDEELPAQRYERDLVQKLKVLRHELSLQQPQAGHCRIEVSREEIFEESYRQIMKMRPKDLKKRLMVKFRGEEGLDYGGVAREWLYLLCHEMLNPYYGLFQYSTDNIYMLQINPDSSINPDHLSYFHFVGRIMGLAVFHGHYINGGFTVPFYKQLLGKPIQLSDLESVDPELHKSLVWILENDITPVLDHTFCVEHNAFGRILQHELKPNGRNVPVTEENKKEYVRLYVNWRFMRGIEAQFLALQKGFNELIPQHLLKPFDQKELELIIGGLDKIDLNDWKSNTRLKHCVADSNIVRWFWQAVETFDEERRARLLQFVTGSTRVPLQGFKALQGSTGAAGPRLFTIHLIDANTDNLPKAHTCFNRIDIPPYESYEKLYEKLLTAVEETCGFAVE

### His-Tag Smurf1 HECT Domain Protein Sequence

MGSSHHHHHHSSGLVPRGSHHIMNHQCQLKEPSQPLPLPSEGSLEDEELPAQRYERDLQKLKVLRHELSLQQPQAGHCRIEVSREEIFEESYRQIMKMRPKDLKKRLMVKFRGEEGLYGGVAREWLYLLCHEMLNPYYGLFQYSTDNIYMLQINPDSSINPDHLSYFHFVGRIMGLVFHGHYINGGFTVPFYKQLLGKPIQLSDLESVDPELHKSLVWILENDITPVLDHTFCVEHNAFGRILQHELKPNGRNVPVTEENKKEYVRLYVNWRFMRGIEAQFLALQKGFNELIPQHLKPFDQKELELIIGGLDKIDLNDWKSNTRLKHCVADSNIVRWFWQAVETFDEERRARLLQ F VTGSTRVPLQGFKALQGSTGAAGPRLFTIHLIDANTDNLPKAHTCFNRIDIPPYESYEKLYEKLLTAVEETCGFAVE

### His-Tagged Smurf1^Trunc^ Protein Sequence

MGSSHHHHHHSSGLVPRGSHHIMNHQCQLKEPSQPLPLPSEGSLEDEELPAQRYERDLVQKLKVLRHELSLQQPQAGHCRIEVSREEIFEESYRQIMKMRPKDLKKRLMVKFRGEEGLDYGGVAREWLYLLCHEMLNPYYGLFQYSTDNIYMLQINPDSSINPDHLSYFHFVGRIMGLAVFHGHYINGGFTVPFYKQLLGKPIQLSDLESVDPELHKSLVWILENDITPVLDHTFCVEHNAFGRILQHELKPNGRNVPVTEENKKEYVRLYVNWRFMRGIEAQFLALQKGFNELIPQHLLKPFDQKELELIIGGLDKIDLNDWKSNTRLKHCVADSNIVRWFWQAVETFDEERRARLLQFVTGSTRVPLQGFKALQGSTGAAGPRLFTIHLIDANTDNLPKAHTCFNRIDIPPYES YEKL YEKLLTAVEETCG

### Smurf1C2 Protein Sequence

MGSSHHHHHHSSGLVPRGSHHIMNHQCQLKEPSQPLPLPSEGSLEDEELPAQRYERDLVQKLKVLRHELSLQQPQAGHCRIEVSREEIFEESYRQIMKMRPKDLKKRLMVKFRGEEGLDYGGVAREWLYLLCHEMLNPYYGLFQYSTDNIYMLQINPDSSINPDHLSYFHFVGRIMGLAVFHGHYINGGFTVPFYKQLLGKPIQLSDLESVDPELHKSLVWILENDITPVLDHTFCVEHNAFGRILQHELKPNGRNVPVTEENKKEYVRLYVNWRFMRGIEAQFLALQKGFNELIPQHLLKPFDQKELELIICGLGKIDVNDWKVNTRLKHCTPDSNIVKWFWKAVEFFDEERRARLLQFVTGSSRVPLQGFKALQGAAGPRLFTIHQIDACTNNLPKAHTCFNRIDIPPYESYEK LYEK LLTAIEETCGFAVE

